# A Toolbox and Crowdsourcing Platform for Automatic Labeling of Independent Components in Electroencephalography (ALICE)

**DOI:** 10.1101/2021.04.06.438576

**Authors:** Gurgen Soghoyan, Alexander Ledovsky, Maxim Nekrashevich, Olga Martynova, Irina Polikanova, Galina Portnova, Anna Rebreikina, Olga Sysoeva, Maxim Sharaev

## Abstract

Independent Component Analysis (ICA) is a conventional approach to exclude non-brain signals such as eye-movements and muscle artifacts from electroencephalography (EEG). Due to other possible EEG contaminations, a rejection of independent components (ICs) is usually performed in semiautomatic mode and requires experts’ involvement. Noteworthy, as also revealed by our study, experts’ opinion about the nature of a component often disagrees highlighting the need to develop a robust and sustainable automatic system for EEG ICs classification. The current article presents a toolbox and crowdsourcing platform for Automatic Labeling of Independent Components in Electroencephalography (ALICE) available via link http://alice.adase.org/. The ALICE toolbox aims to build a sustainable algorithm not only to remove artifacts but also to find specific patterns in EEG signals using ICA decomposition based on accumulated experts’ knowledge.

The difference from previous toolboxes is that the ALICE project will accumulate different benchmarks based on crowdsourced visual labeling of ICs collected from publicly available and in-house EEG recordings. The choice of labeling is based on estimation of IC time-series, IC amplitude topography and spectral power distribution. The platform allows supervised ML model training and re-training on available data subsamples for better performance in specific tasks (i.e. movement artifact detection in healthy or autistic children). Also, current research implements the novel strategy for consentient labeling of ICs by several experts. The provided baseline model shows that it can be used not only for detection of noisy IC but also for automatic identifications of components related to the functional brain oscillations such as alpha and mu-rhythm.

The ALICE project implies the creation and constant replenishment of the IC database, which will be used for continuous improvement of ML algorithms for automatic labeling and extraction of non-brain signals from EEG. The toolbox and current dataset are open-source and freely available to the researcher community.

## INTRODUCTION

Electroencephalography (EEG) signal reflects bioelectrical activity of brain neuronal networks. For more than a century human neuroscience and clinical research applied scalp EEG recording to study and assess a wide scope of sensory and cognitive functions. One of crucial steps of EEG preprocessing is ‘purifying’ the EEG signal by an extraction of electrical activity of non-neuronal origin such as eye-movement and muscle artifacts. For the recent decades Independent Component Analysis (ICA) offered a solution to this problem based on isolation of statistically independent sources called independent components (ICs) as linear combinations of signals from electrodes [1,2]. A source of each IC can be projected onto electrode cap and estimated via timecourse and spectral power. For example ICA allows to identify components related to eye-movement and muscle artifacts based on the specific characteristics their bioelectrical signals, e.g. frequency and spatial distribution [3,4]. However, due to other frequent contaminations of EEG, a rejection of non-brain ICs is usually performed in semiautomatic mode under visual inspection of researchers. Herewith, a labeling of ICs by different experts, can substantially disagree, which might considerably affect the further analysis and reproducibility of EEG results [5]. Artifact rejection by ICA in children and patient EEG is especially challenging even for experts. The dependence of EEG analysis from subjective opinions of experts may explain that EEG data have been rarely included in large-scale studies or meta-analyses. For this reason the automatic algorithms for EEG processing are the main objectives of many research groups [6–11].

To create a robust and sustainable automatic system for EEG ICs classification, one needs to extract most informative features from ICs as well as to have an appropriate machine learning (ML) model inside the system. The objective labeling of ICs is the crucial step in training and validating this model. Training of ML algorithms to automatically identify artifactual ICs will allow to set up more objective methodology for EEG pre-processing.

Currently, there is a limited number of projects aimed at creating an automatic cleaning system of the EEG signal. For example, Automatic EEG artifact Detection based on the Joint Use of Spatial and Temporal features (ADJUST) [8] and Fully Automated Statistical Thresholding for EEG artifact Rejection (FASTER) [9], which are empirical threshold-based algorithms. Machine learning approach introduced in Multiple Artifact Rejection Algorithm (MARA) [10] and algorithms from the studies of Frølich and colleagues [4] and Tamburro and colleagues [11]. SASICA software [3] is an EEGLAB Matlab plug-in [3], includes ADJUST, MARA, FASTER and some other methods. All these studies used their own private datasets for training and validation purposes. Those datasets were relatively small, consisting of several hundred ICs. In most cases, each IC was annotated by only one expert. These problems complicate estimation of algorithms’ actual performance and their comparison. Moreover, the lack of a large dataset with verified annotation limits machine learning models potential performance.

Pion-Tonachini and colleagues [12] made a step forward solving this problem proposing ICLabel Toolbox, which includes the annotation tool with crowdsourcing mechanics, datasets and several machine learning algorithms. The annotation tool provides an interface to label a particular IC from the database by visualizing different components characteristics. In this toolbox the ML algorithms are based on artificial neural networks and claimed to be the fastest and most accurate as compared with other studies.

While the ICLabel project is a great resource for automatic artefact rejection in EEG, it has several drawbacks. The first one is potentially poor annotation quality as ICs can be annotated by a non-expert user. It means that even if an ML algorithm has a high accuracy, the predicted classes may be wrong as ICs have no order or intrinsic interpretations and their classification by experts requires practice. Potential technical issues that prevent best performance from experts are inability to see other ICs from the same EEG record which is useful in ambiguous cases (e.g. horizontal eyes component can be projected into two ICs and to see them in parallel helps to infer its nature) and limitation of component time series plots to only 3 seconds ranges. Noteworthy, clinical experts usually require at least 30 seconds to properly detect various slow wave components or for example alpha rhythm which can be weakly expressed in a short time interval. Other limitation of ICLabel that was pointed by the authors themselves is limited variety of EEG data, as their dataset does not contain data from infants and most clinical groups.

The current article presents a toolbox and crowdsourcing platform for Automatic Labeling of Independent Components in Electroencephalography (ALICE) which is available via link http://alice.adase.org/. The ALICE toolbox aims to build a sustainable algorithm to remove artifacts and find specific patterns in EEG signals using ICA decomposition and to account for major drawbacks of previous approaches.

To develop a sustainable ML-based EEG components classification system, it is necessary to have two components: a high-quality labelled dataset and a proper machine learning pipeline to train and validate models.

Thus, the first aim of the ALICE project is to create a high-quality dataset with IC labels. In order to achieve this goal, we need:

- to define a rigorous set of possible IC classes that would cover high variety of cases and be easily understandable by experts
- to annotate IC reliability by combining opinions from multiple experts
- to resolve possible bad concordance between experts by various merging strategies
- to attract researchers to share their datasets, including unique EEG recordings from rare clinical groups.

The second aim of the ALICE project is to develop a robust, but flexible machine learning pipeline for the automated IC classification. The machine learning module includes an implementation of various features (both well established and original), multiple ML models and the validation pipeline. The community is also invited to develop their own models using our dataset which is available via link http://alice.adase.org/downloads.

The other ambitious goal for ALICE development is automatic identification of components related to the functional brain oscillations such as alpha and mu-rhythm. Mu rhythm overlaps with alpha rhythm in a frequency range of 8−13 Hz but has different form of oscillations and localization at scalp electrodes. While alpha rhythm is recorded predominantly from the occipital lobes with closed eyes and suppressed when eyes open [13], mu rhythm emerges over the sensorimotor cortex and attenuated during movements. Importantly, mu rhythm does not react to opening or closing the eyes [14]. Despite the described differences, the automatic separation of mu from alpha waves in EEG is challenging and drawing attention of many methodological studies [15,16]. Still, the identification of mu rhythm often requires visual inspection and expertise. The ALICE toolbox aims to accumulate expert labeling of alpha and mu rhythms to improve automatic identification of functional brain oscillations by supervised machine learning.

## MATERIALS AND METHODS

### ALICE toolbox high-level architecture

ALICE contains two modules (Fig 1):

**Fig 1.**
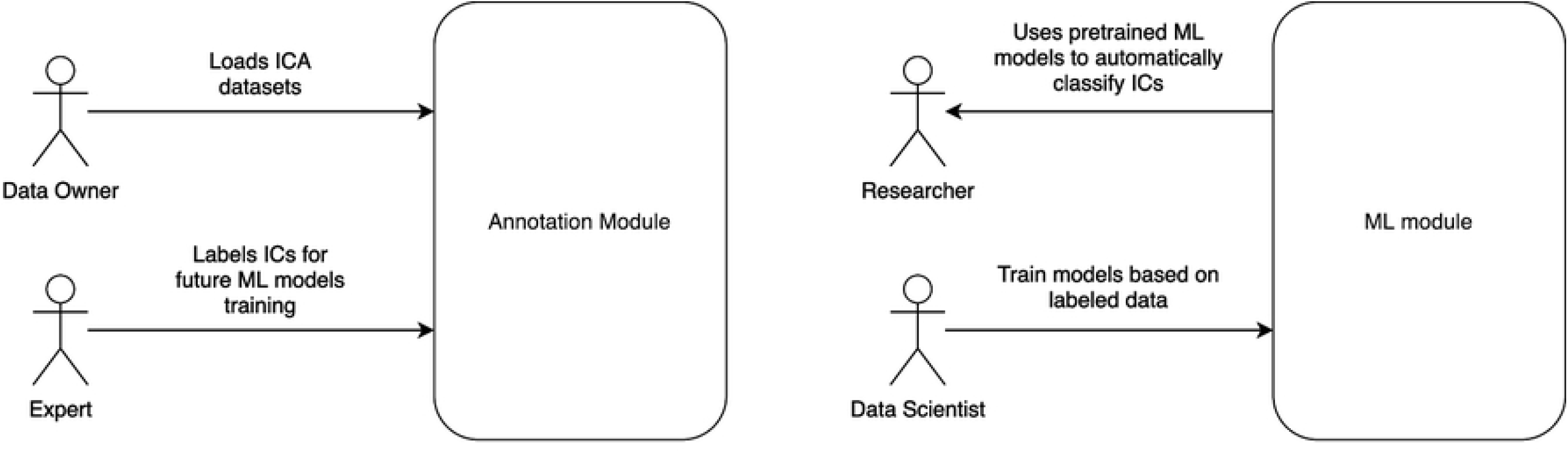
ALICE toolbox high-level architecture and user roles. Annotation module UI serves for Data Owners to upload IC data to the database and for Experts to provide annotation on existing or newly uploaded data. Data Scientists and Researchers work with ML module: the former train models based on selected samples from the database, the latter take pretrained models to work with their own data (online or offline). Online version of the ALICE toolbox is available at http://alice.adase.org/

- Annotation module, which consists of a web-based user-interface (UI) and a database. An HTTP API allows uploading ICs data to the database. Web-based UI allows experts to label uploaded data for future ML models training and validation
- ML module based on a Python library which allows to train ML models based on expert annotations and use pre-trained ML models to apply to new IC data.

### Annotation module

By annotation we mean a process of manual IC labeling by experts based on various data visualization tools available at ALICE platform, such as component topomap, plots of timeseries and power spectrum. An expert may choose IC labels from a predefined number of options.

We propose a set of IC component labels including major artifact types with subtypes as well as brain signal subtypes:

- Eyes artifacts
  - Eyes artifacts of any type.
  - Horizontal eye movements. Components that represent activity during eye movements in horizontal directions.
  - Vertical eye movements. Components that represent activity during eye movements in vertical directions.
- Noise
  - Line noise. Line current noise that is evoked by surrounding electrical devices.
  - Channel noise. The noise associated with channels that can be marked as bad.
- Brain activity
  - Brain activity of any type.
  - Alpha activity.
  - Mu activity.
- Muscle Activity. Artifacts from a recording of muscle activity on the head surface.
- Heartbeat artifacts. Artifacts that represent electrocardiographic activity.
- Other. Components with clear nature which label is not listed in labeling system, for example it could be breathing. Experts could comment the label for them in the comments section. The ALICE developers collect data from comments and based on them expand the list of labels in the posterior versions of tool.
- Uncertain. Components with unclear nature,

Annotation process is supported by the web-based UI, see Fig. 2. An expert has the following data visualization options:

**Fig 2.**
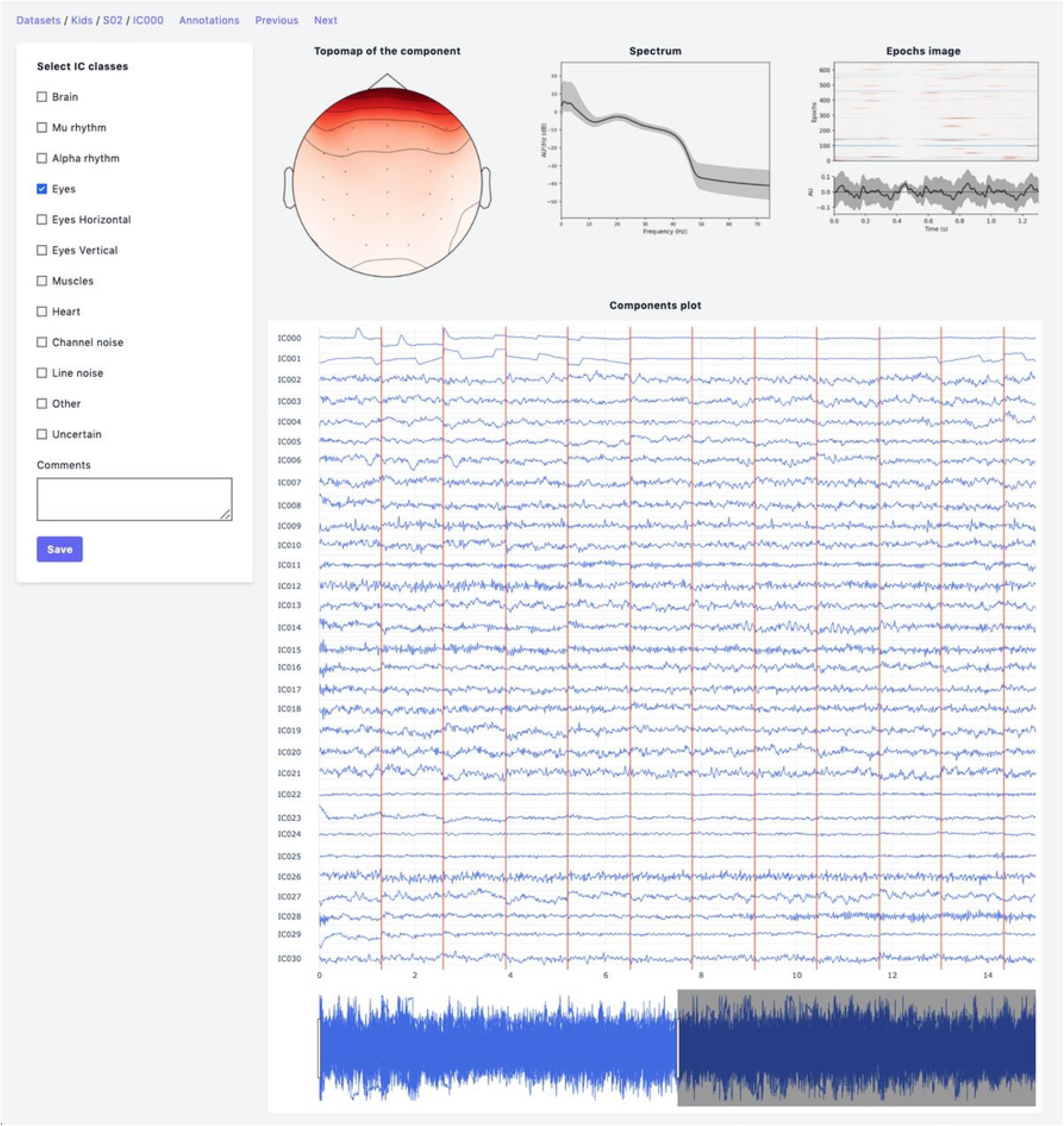
Annotation Module interface. Top row: IC topomap and spectrum as well as Epochs image, illustrating the color-coded amplitude fluctuations (arbitrary units) of the IC in all available epochs (time relative to sound presentation on the x-axis, epochs on the y-axis). Image at the bottom shows averaged values of ICs timeseries; bottom row: all IC for considered subject are plotted together.

- Topomap of IC weights.
- Power spectrum density plot.
- Plot of all ICs time series for the current subject. Time-series length is 30 seconds with the possibility of scrolling and zooming selected time interval.
- Epoch image illustrating the color-coded amplitude fluctuations of the IC in all available Epochs together with averaged values of ICs timeseries. This plot is useful for annotation of epoched data.

After the labeling process by a particular expert is finished, the data of ICs with annotations can be packed to an archive by the annotation module administrator. Then, annotated data becomes available at the Downloads page of ALICE toolbox and could be used both by the experts and ALICE data scientists.

### Machine learning pipeline

There could be many discrepancies between experts’ annotations due to ambiguities in IC patterns, data quality and expert level. This means that we need to create final IC labels in the dataset as a function of the individual annotations. So, before conventional ML pipeline steps, such as *Features calculation* and *ML models training and selection*, we need to add an additional one - *Data labels aggregation* step. The whole data processing and machine learning pipeline is presented in Fig 3 and each step is discussed in detail below.

**Fig 3.**
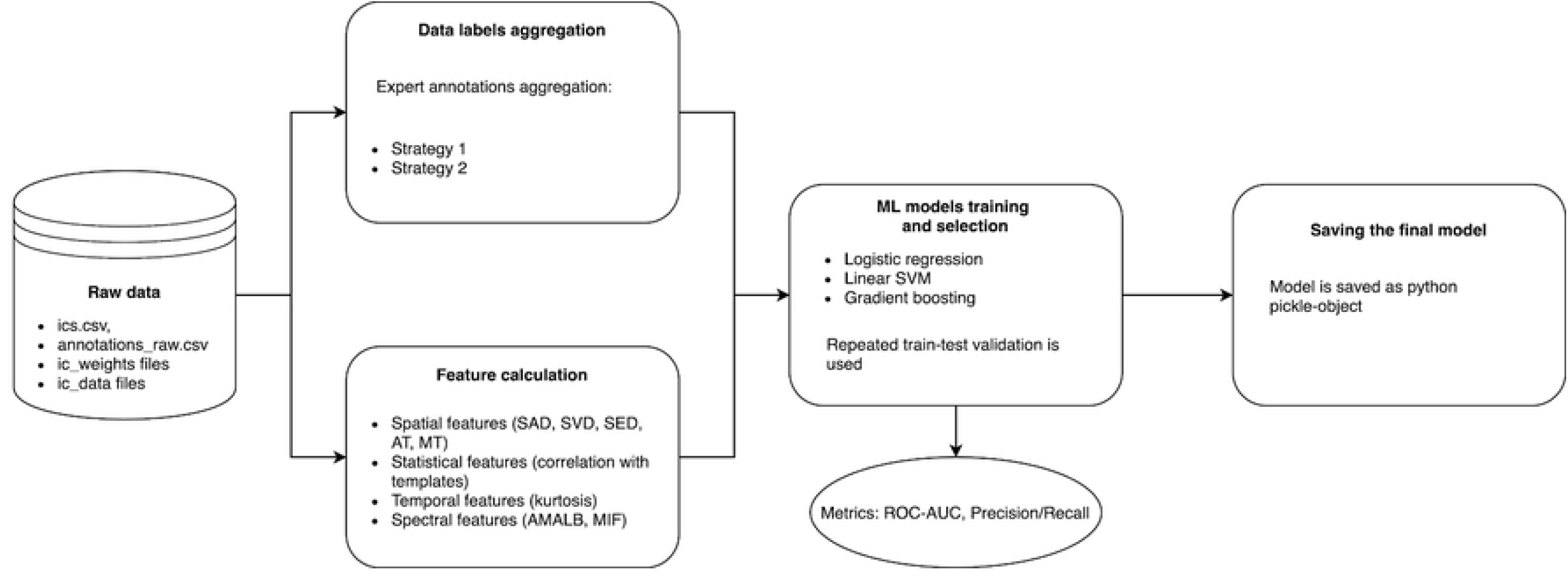
Data processing and machine learning pipeline in ALICE. Raw ICs data with annotations are passed to Data labels aggregation and Feature calculation blocks to form a labelled dataset and extract informative features from ICs. Three ML models are trained with repeated train-test validation and different model quality metrics are calculated. The best model is then selected and could be exported as a Python pickle-object.

#### Data labels aggregation

This part aims to create a boolean variable between each component and each IC class that represents whether the certain activity is present or not in a particular component. First step is to create an annotation table (Fig 4A). *Annotation* is a term denoting the labeling produced by an expert to a particular component. Each expert has their own unique opinion about the component’s ICA class. Our goal is to develop an approach to grouping expert annotations to form a common opinion on each component.

**Fig 4.**
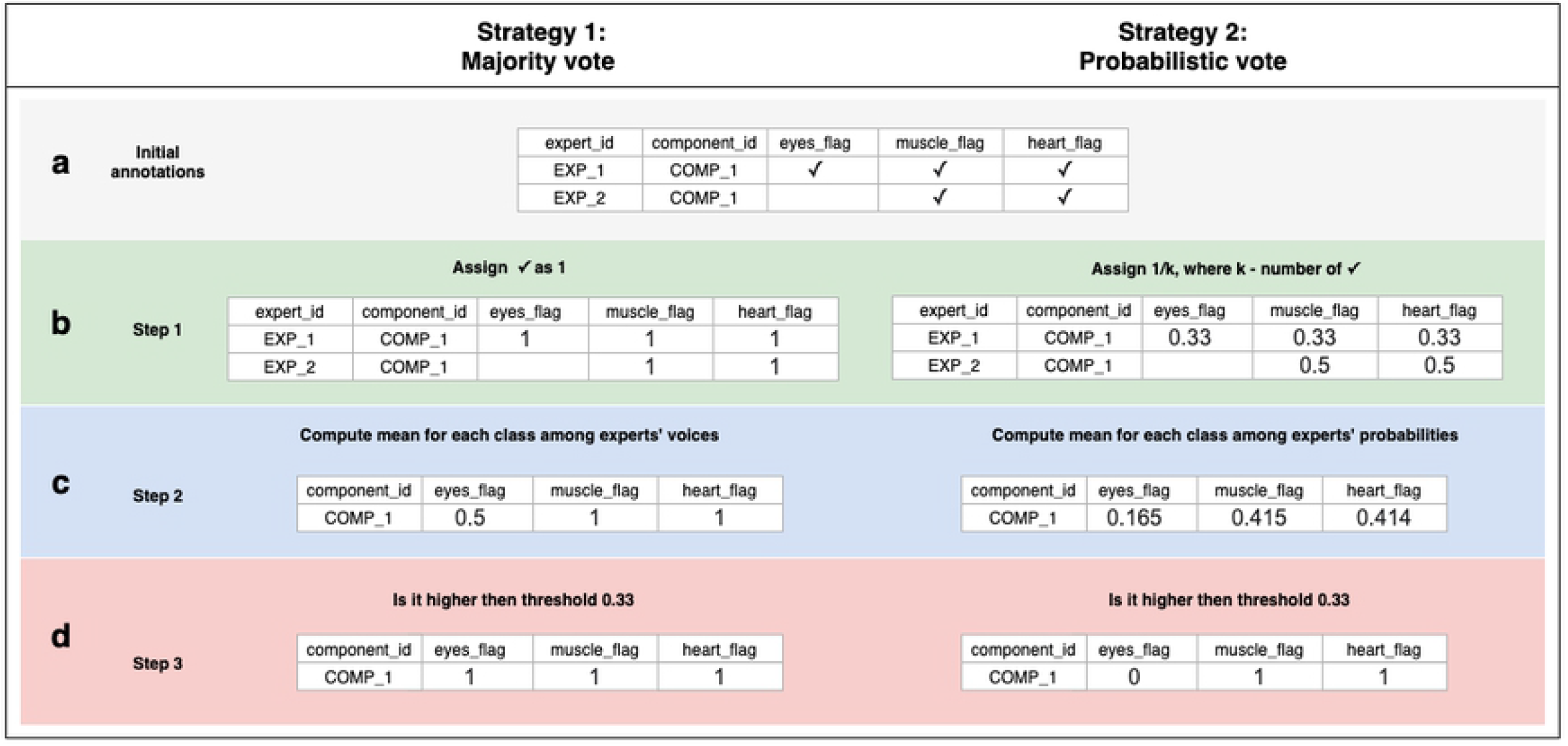
Data labeling strategies. There are several annotations for one component from various experts, but we strive to designate its belonging to a particular class strategies of data belling aggregation, those are “Majority vote” and “Probabilistic vote”. (A) - is a table of annotations of a specific component. (B) - transform into a matrix of *probabilities* that each component belongs to a particular class. (C) - group the experts’ opinions using the mean of the *probabilities* to obtain a *weighted probability*. (D) - determine whether the weighted probability is higher than the threshold or not

A simple voting strategy seems to be a logically correct option: if the majority of experts choose that a component contains a particular activity, for example, an eyes artefact, then this component is classified as an eyes’ artefact. This approach is the basis of Strategy 1 which is called *“Majority vote”*, although it does not require that the majority (more than 50%) of the experts assign the component this particular label. The threshold can be varied and requires, for example, more than 33% of experts to choose uniformly. In other words, by grouping experts’ annotations, we form the average of the experts’ votes (Fig 4C). We will consider this average value as the *probability* of assigning the component to a specific class. If the probability is higher than the threshold, we assume that the component encodes the given IC class; otherwise, it does not (Fig 4D).

Nevertheless, if an expert assigned a component to several classes, it means s/he recognizes that several types of activity are mixed in it. This situation can lead to ambiguous results if the expert acted with an approach where s/he labels mixed components with all types of activity s/he believes are potentially intermixed in a particular IC. If we were to use Strategy 1 for such situations, it would lead to low quality of the target variable as IC with only one label is more genuine representation of this class than the component that contains mixture of the artificial and brain activity. An example of what this can affect is illustrated in Fig 5. We see that the component, due to such markup, is assigned to all classes simultaneously.

**Fig 5.**
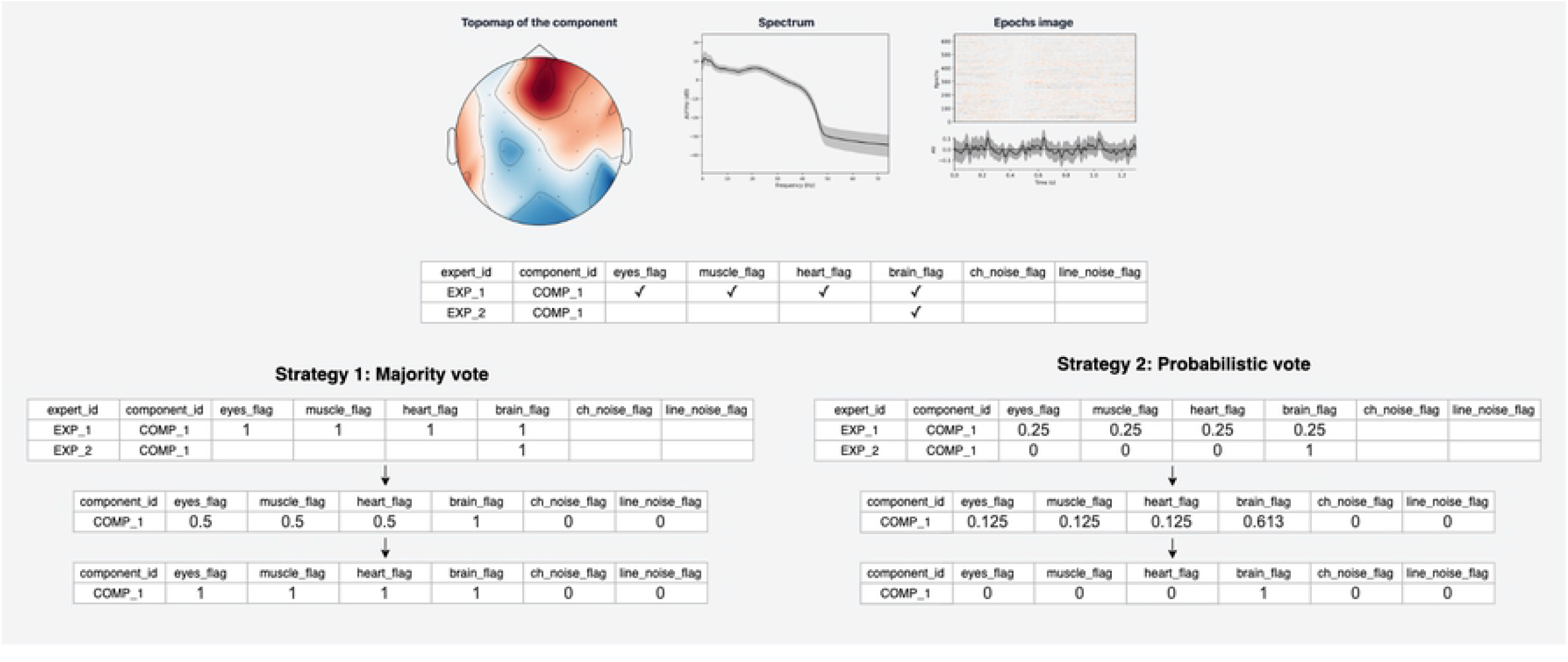
Discrepancies in experts’ opinions and difference in Strategies handling. Many IC components consist of more than one activity type. Experts had different approaches in labeling samples with mixed activity. Expert 1 labelled all activity types that are present in the component. Expert 2 annotated components only with a clear pattern of chosen IC class. Within a provided example we can see that the application of “Majority vote” will assign this component as target for many classes, that can affect our model performance. On the other hand, Strategy 2 handle such situation in more appropriate way.

In order to overcome this situation, Strategy 2 was developed and titled as “Probabilistic vote”. Imagine that, when labeling a component, an expert has one vote, which they equally distribute among all the classes to which they attributed this component. In other words, if a person marks a component as eyes and as muscles and as heart, then with a *probability* of 0.33, they assign it to each of these classes (Fig 4B). Further, these *probabilities* are again averaged (Fig 4C). Then, again, a threshold is chosen, according to which it is decided whether this *weighted probability* will be transformed to 1 or 0 (Fig 4D). The threshold of 0.33 was chosen as the optimal threshold for the current data, assuming that components that consists of three or less labels still represent the genuine pattern of interest for the model. This approach is rather useful in cases where mixed nature of component can affect target variable and example for that is provided in Fig 5C.

The threshold value is highly dependent on the level of agreement between experts since a too tight threshold with a low agreement will significantly reduce the number of objects. On the other hand, a weak threshold with a high agreement will lead to noisy, ambiguous components in the training set. We decided to use equal threshold of 0.33 for both strategies. The threshold change for Strategy 1 will make sense with an increase in number of experts.

#### Agreement between experts

We also computed metrics of expert agreement to be able to compare annotation quality of various classes as well as datasets. For the case of two experts we propose using Kohen’s kappa [17].

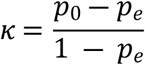

where *p*_0_ is the relative observed agreement (similar to accuracy), *p*_*e*_ is the hypothetical probability of agreement by chance.

For the case of multiple experts we propose using Fleiss’ kappa [18] which has a similar formula with a different definition of *p*_0_ and *p*_*e*_.

Based on the metrics from [12] we computed the inter-expert correlation between experts to compare our level of agreement with level of agreement in ICLabel.

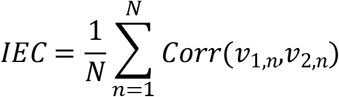

where *N* – number of components that were marked by both experts, *v*_1,*n*_ *–* annotation vector made by 1^st^ expert corresponding to *n*^*th*^ component, *v*_1,*n*_ *–* annotation vector made by 2^nd^ expert corresponding to *n*^*th*^component

#### Features calculation

In order to reduce data dimensionality while preserving most characteristic information for each IC class we calculate specific temporal and spatial features of each signal. Most features are well established and based on previous research. Still, we introduced some modifications to existing ones and treated them as new features.

Among the established features are:

- Kurtosis of the component time series [8–11]. By definition, kurtosis is the fourth standardized moment of the time series. In case of epoched data we calculate an average of the feature computed for each epoch separately. It helps to distinguish ICs that correspond to eyes and brain activity.
- Maximum Epoch Variance [8,11] is used to detect eye movements. The value of this feature is a ratio between the maximum of the signal variance over epochs and the mean signal variance. As proposed in [8], when calculating this feature, we excluded one percent of the largest variance values to improve its robustness.
- Spatial Average Difference (SAD), Spatial Variance Difference (SVD) and Spatial Eye Difference (SED). Spatial features proposed in [8] depend on IC weights of eyes-related electrodes. SAD is calculated as the difference between channel weight averages over frontal and posterior regions. SVD is the difference between weight variances in these regions. These are used to distinguish vertical eye movements. SED is the difference between the absolute values of weight averages in the left eye and right areas. This feature detects horizontal eye movements.
- Myogenic identification feature (MIF) [11] is used to detect muscle activity and is calculated as relative strength of the signal in the 20 to 100 Hz band.
- Correlations with manually selected signal patterns [11]. We use these to detect eye blinks and eye movements.

ALICE is, to the best of our knowledge, the first tool with the possibility of Mu and Alpha rhythms annotation and classification. Thus, some features must be specific to these components’ spatial and temporal properties.

It is well known that Alpha activity is localized in occipital and central areas with increased power in 6-12 Hz. On the other hand, Mu rhythm is generated in central and frontal areas with increased power in the same frequency band. By the topography related features, we used those electrodes that maximally emphasize the contrast between mu and alpha localization. Thus, the original features include:

- Mu Topography (MT). A feature which is sensitive to topomaps of mu rhythm ICs, where *Mu is* the following set of electrodes in 10-20: ‘Fp1’, ‘Fpz’, ‘Fp2’, ‘F3’, ‘Fz’, ‘F4’, ‘Fc3’, ‘Fcz’, ‘Fc4’, ‘C3’, ‘Cz’, ‘C4’

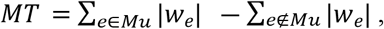
- Alpha Topography (AT). A feature which is sensitive to topomaps of alpha rhythm ICs where *A is* the following set of electrodes in 10-20: ‘C3’, ‘Cz’, ‘C4’, ‘Cp3’, ‘Cpz’, ‘Cp4’, ‘P3’, ‘Pz’, ‘P4’, ‘O1’, ‘Oz’, ‘O2’

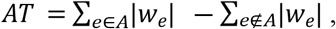
- Average Magnitude in Alpha Band (AMALB). The ratio between average amplitude in alpha band (6-12Hz) and average amplitude in other frequencies (0-6 Hz; 13-125 Hz) which is sensitive to alpha ICs.

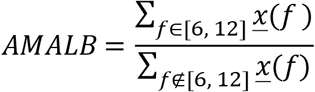

#### ML models training and selection

The current version of ALICE Toolbox provides three different machine learning models: logistic regression (LR), linear support vector machine (SVM) and gradient boosting (XGB). These models are built on different principles and are rather simple compared to neural networks and deep neural networks. Keeping in mind a relatively small initial dataset we considered the aforementioned three models as an optimal initial model choice. All of them are optionally available for new training and testing procedures in ALICE. In particular, we used the LR implementation from scikit-learn package [19] with default parameters (including regularization parameter C=1.0, L2 penalty and liblinear solver). Linear SVM is taken from scikit-learn package [19] with default parameters (including regularization parameter C=1.0). Finally, we used the XGB model implementation from XGBoost package [20] with default parameters max_depth=4 and 30 estimators.

In ALICE we implement repeated train-test split cross-validation technique. We trained the model on 70% of samples and validated on the rest 30% and did not optimize any hyperparameters on cross-validation. We performed this procedure 50 times using different random train-test splits estimating two main metrics of classification accuracy: Area Under the Receiver Operating Characteristic Curve (ROC-AUC) and F1-score using the implementation of scikit-learn package [19]. ROC-AUC was used as an overall metric of a model performance for different thresholds and was considered as the main one. F1 was used as a performance metric of model optimal splits and was considered as an additional metric.

#### Initial dataset

The ALICE project aims to involve the neurophysiological community in labeling existing publicly available and new IC datasets to improve ML models’ quality. However, the Baseline model trained on the dataset provided by IHNA&NPh RAS is already available to users.

EEG data were recorded using the NeuroTravel system with sampling rate 500 Hz and with 31-scalp electrodes arranged according to the international 10–10 system and included the following electrodes: ‘Fp1’, ‘Fpz’, ‘Fp2’, ‘F3’, ‘Fz’, ‘F4’, ‘F7’, ‘F8’, ‘FC3’, ‘FCz’, ‘FC4’, ‘FT7’, ‘FT8’, ‘C3’, ‘Cz’, ‘C4’, ‘CP3’, ‘CPz’, ‘CP4’, ‘P3’, ‘Pz’, ‘P4’, ‘TP8’, ‘TP7’, ‘T3’, ‘T4’, ‘T5’, ‘T6’, ‘O1’, ‘Oz’, ‘O2’. Ear lobe electrodes were used as reference, and the grounding electrode was placed centrally. The initial dataset consists of recordings from 20 typically developing children aged 5 to 14 years. Within the experiment’s framework, sound stimulation was performed according to the odd-ball paradigm with a standard stimulus of 1000 Hz and two deviant stimuli at 980 Hz and 1020 Hz. The interstimulus interval was 400 ms. Stimulus intensity were 75 dB.

Obtained data were filtered (0.1 - 40 Hz) and divided into epochs (−500; 800 s), where noisy epochs were removed by threshold (350 mV). Only the first 650 epochs were used for posterior ICA decomposition (FASTICA) with resampling on the level of 250 Hz. Final data that were uploaded into ALICE consisted of 30 ICA components. All preprocessing steps were done using the MNE Python package [21].

The data annotation for training the Baseline model was carried out by two experts - experienced scientists of the Institute of Higher Nervous Activity and Neurophysiology of RAS. For the correct work with the program, they received an instruction, which outlined the main steps they took when working with ALICE. Experts’ main task was set as follows - to mark each component using the set of labels: Eyes, Horizontal eye movements, Vertical eye movements, Line noise, Channel noise, Brain, Alpha activity, Mu activity, Muscle activity, Other, Uncertain. Following the instructions, if an expert saw that a component consisted of several activity types, s/he can assign the component to several classes. For example, among annotated components there were often components that were marked both as eye artifacts and muscle activity simultaneously.

#### Ethical statement

The dataset was obtained from the research project conducted according to the guidelines of the Declaration of Helsinki and approved by the Ethics Committee of the Institute of Higher Nervous Activity and Neurophysiology (protocol code 3, date of approval 10 July 2020). All children provided their verbal consent to participate in the study and were informed about their right to withdraw from the study at any time during the testing. Written informed consent was also obtained from a parent/guardian of each child.

## RESULTS

*Data labeling aggregation*. First, we explored the level of consistency between two annotators for various IC classes. Due to limited available data and only two annotators we decided to merge some classes with a small number of label matches between annotators. One reason for this small number could be the possible difference in labeling strategies between the experts, as was discussed in the Methods section. The final manipulations with class labels are:

- Eyes, Horizontal eyes movement, Vertical eyes movement were merged to the one Eyes class
- Line noise labels were dropped due to lack of actual line noise in available data

For the rest of the IC classes we used the following aggregation strategies based on the total number of positive samples of each class (see Table 1). When the samples of a particular class were weakly represented, we took Strategy 1 to have enough labelled samples for the model fitting, otherwise we took Strategy 2. The details of Strategy 1 and Strategy 2 are explained in the Methods section.

**Table 1.** IC classes and corresponding aggregation strategies based on the total number of positive samples of each class.

| IC Class | Brain activity | Alpha brain activity | Mu brain activity | Eyes | Muscles | Heart | Channel noise |
| --- | --- | --- | --- | --- | --- | --- | --- |
| **Strategy** | 2 | 1 | 1 | 2 | 2 | 1 | 2 |

The final number of positive labels and concordance between two experts are shown in Table 2.

**Table 2.** Number of samples and Kohen’s kappa for each class. The total number of samples was 620.

| Label | Number of samples | Cohen's kappa |
| --- | --- | --- |
| Brain | 449 | 0.47 |
| Alpha | 60 | 0.13 |
| Mu | 92 | 0.22 |
| Eyes | 78 | 0.10 |
| Muscles | 135 | 0.36 |
| Heart | 231 | 0.04 |
| Channel noise | 48 | 0.12 |

According to arbitrary settled thresholds [22], the agreement between two experts’ opinions was highest but still moderate (<0.4) only for labeling the ICs of brain signals. The other ICs were labeled with rather poor agreement between experts (Table 2). The Inter-expert correlation between our experts equals 0.43 and approximate level of agreement was also reviewed in [12]. Based on the experts’ comments, we understood that many of the IC components contain more than one activity type. This mixture led to uncertainty for experts’ labeling strategy. Summing up their annotations and based on the comments, we can conclude that one expert was inclined to label only those components where a clear pattern of chosen IC class could be detected. Another expert labelled all activity types that are present at a given components, even when there was only slight indication of its presence in a clearly multi-nature ICs. This difference in labeling strategies produced rather poor agreement even for (usually well recognized) Eyes activity. The annotation dataset available via http://alice.adase.org/downloads and [23].

### IC classification

As it can be seen from Table 2, many classes are relatively small. This leads to imbalanced classification tasks for example for Alpha, Mu and Channel noise IC classes. In this case Precision-Recall (PR) curve better reflects classifier performance compared to conventional ROC-curve. So, we explored LR, XGB and SVM as ML models and calculated both ROC-AUC and PR-AUC scores as performance measures. All the models showed close performance for most ICs classes (see Fig.6 for ROC curves and Table 3 for values) based on ROC-curves. Brain, Eyes and Muscles models showed the best performance among others with ROC-AUC greater than 0.9. We were not able to train a good model for Heart ICs detection due to poor labeling as suggested by the lack of consistency among experts and probably not very specific extracted features.

**Table 3.** Average ROC-AUC values and their standard deviations.

| ROC-AUC | Brain | Alpha | Mu | Eyes | Muscles | Heart | Channel noise |
| --- | --- | --- | --- | --- | --- | --- | --- |
| LR | 0.93<br>(0.02) | 0.83<br>(0.05) | 0.83<br>(0.03) | 0.91<br>(0.03) | 0.89<br>(0.03) | 0.64<br>(0.04) | 0.81<br>(0.05) |
| XGB | 0.92<br>(0.02) | 0.81<br>(0.05) | 0.83<br>(0.03) | 0.89<br>(0.04) | 0.88<br>(0.02) | 0.61<br>(0.03) | 0.77<br>(0.05) |
| SVM | 0.93<br>(0.02) | 0.81<br>(0.05) | 0.82<br>(0.03) | 0.92<br>(0.03) | 0.90<br>(0.03) | 0.55<br>(0.10) | 0.61<br>(0.09) |

**Fig 6.**
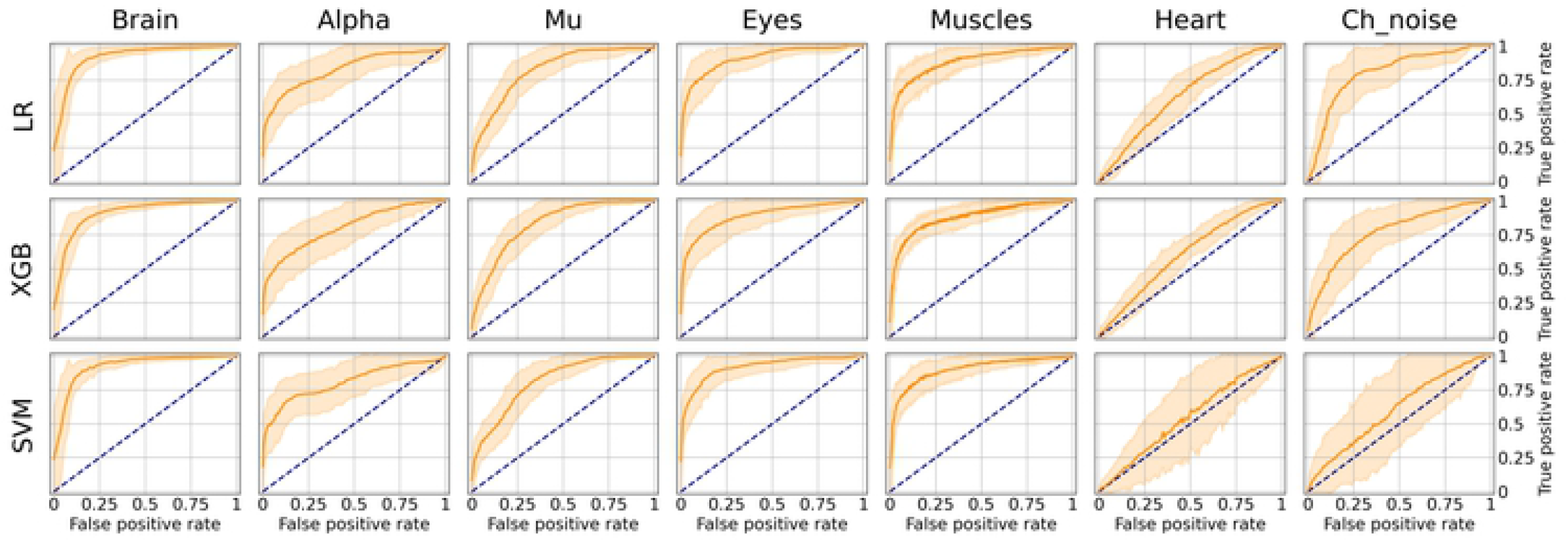
Aggregated Receiver Operating Characteristic (ROC) curves for all IC types and ML models. The solid line indicates the mean curve and the colored area indicates the 95% confidence interval for the ROC curve. The best classification results were achieved for the Brain Muscles and Eyes ICs. For the Alpha Mu and Channel Noise classes, the scores are also high, however the stability is lower, especially in the case of detecting Channel Noise using SVM. Finally, the performance on Heart components was poor, which could be due to low expert concordance. The blue line on each plot represents the no-skill classifier which assigns labels at random. Thus, we can consider the performance of a particular model on a particular label type statistically significant, if the confidence interval lies above the blue line. Thus, most of our models classify the components significantly better than at chance, expert for SVM that was not able to do it for Heart and Channel noise component.

However, the picture is a bit different when analyzing PR-curves and PR-AUC values (see Figure 7 and Table 4). As we mentioned, PR curves better indicate classification performance in case of imbalanced data, which results in worse performance for Alpha, Mu and Eyes IC types, all of which have fewer positive labels compared to Brain or Muscles IC classes. It also can be seen that for Heart and Channel Noise classes all of the models and SVM in particular performed poorly. The possible reasons for this might be both a small number of samples in each class and a low level of agreement between the annotators resulting in poor labeling quality and lack of robustness. Probably, new strong predictive features should be developed to address these types of artifacts. We also provided F1-score values (see Table 5), that is alternative statistics based on precision-recall interaction. The need of further investigation of the models’ performance on Heart and Channel Noise IC classes is also backed up by the low F1-score which for these classes is significantly lower compared with the rest IC types.

**Table 4.** Average PR-AUC values and their standard deviations.

| PR-AUC | Brain | Alpha | Mu | Eyes | Muscles | Heart | Channel noise |
| --- | --- | --- | --- | --- | --- | --- | --- |
| LR | 0.96<br>(0.02) | 0.59<br>(0.08) | 0.50<br>(0.07) | 0.74<br>(0.06) | 0.77<br>(0.05) | 0.46<br>(0.04) | 0.23<br>(0.06) |
| XGB | 0.96<br>(0.01) | 0.54<br>(0.10) | 0.48<br>(0.07) | 0.71<br>(0.07) | 0.75<br>(0.05) | 0.45<br>(0.03) | 0.27<br>(0.09) |
| SVM | 0.96<br>(0.02) | 0.59<br>(0.08) | 0.49<br>(0.07) | 0.76<br>(0.06) | 0.79<br>(0.05) | 0.41<br>(0.07) | 0.13<br>(0.04) |

**Table 5.**
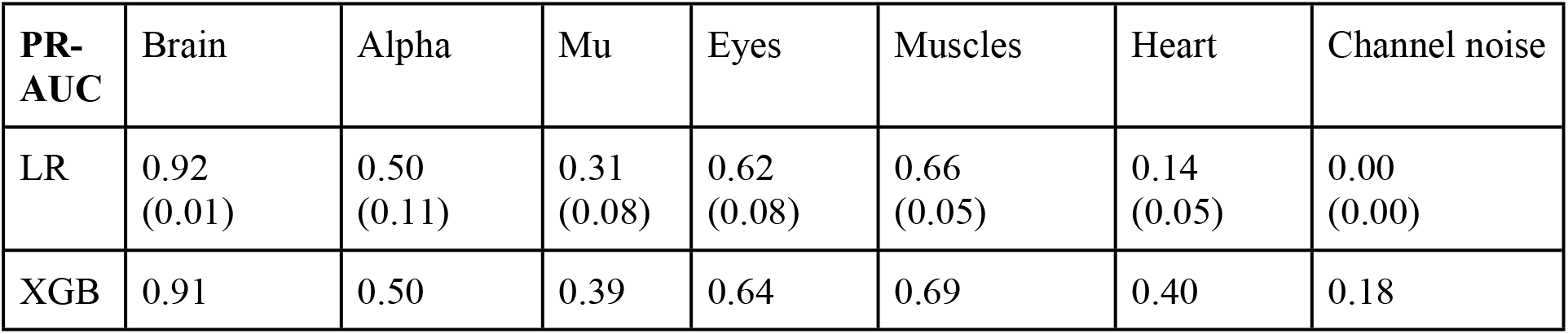
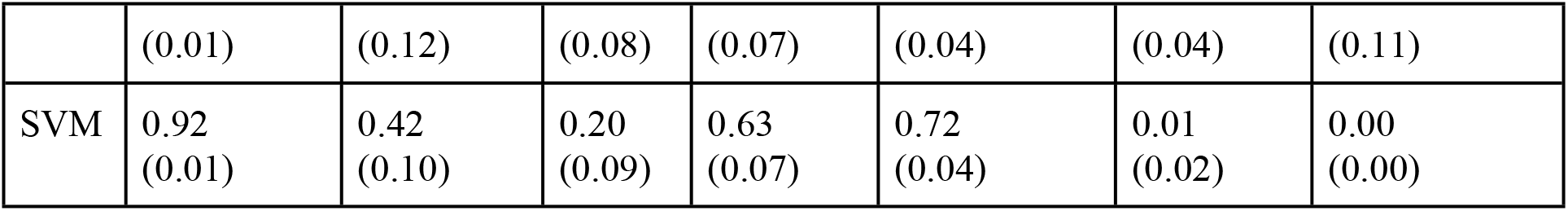
Average F1-scores and their standard deviations.

**Fig 7.**
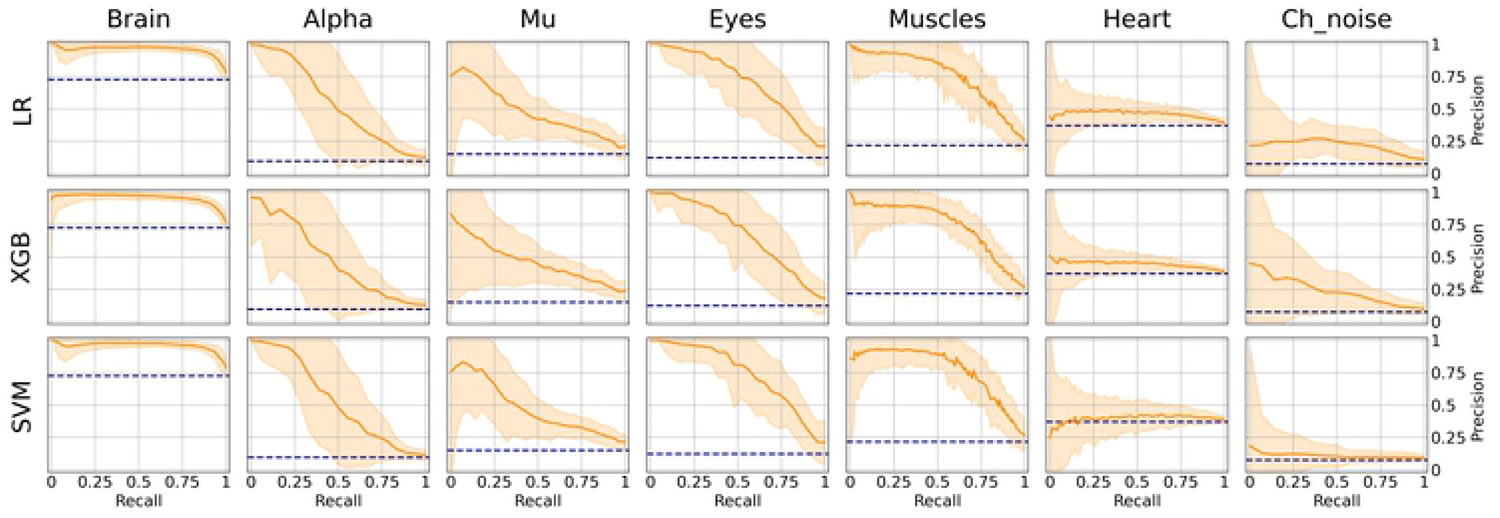
Aggregated Precision Recall (PR) curves for all IC types and ML models. The solid line indicates the mean curve and the colored area indicates the 95% confidence interval for the PR curve. PR curves better indicate classification performance in case of imbalanced data, which can be seen in worse results for Alpha, Mu, Eyes and especially Channel Noise IC types, all of which have fewer positive labels compared to Brain or Muscles IC classes. As with the ROC curves, we can claim that on all IC types except for Heart and Channel Noise, our models perform significantly better than the unskilled classifier.

It is worth mentioning, that the main reason for measuring PR-AUC was to compare the performance of the models with each other. In general, specific PR-AUC values, unlike ROC-AUC, do not reflect the model’s performance. For that, it is better to refer to the PR-curve itself. Each point on this curve corresponds to certain levels of precision and recall, that are closely related to type I and II errors respectively. This can be achieved by choosing the right threshold (by default, each model predicts probabilities for each class that can be interpreted as either True or False by comparing with the threshold value). To better illustrate this idea, consider the following example. Suppose, we want to detect muscles with the recall of 0.75 (that is, we will detect 75% ICs with muscular activities). Then, by looking at Figure 7, we can see that SVM will achieve precision value of about 0.7, which means that out of all ICs that were selected, about 70% will correspond to Muscles.

We chose an ML model for each IC type based on the ROC-AUC score if the class is fairly balanced (Brain and Heart and Muscles) and based on PR-AUC if the class is unbalanced. Thus, we selected PR for Brain, Alpha Mu and Heart, XGB for Channel Noise and SVM for Eyes and Muscles.

## DISCUSSION

ICA is a powerful tool for the segregation of various type of activities from the raw EEG data. It is widely used for detection of different artifacts such as eye blinks or muscle contractions. Nevertheless, IC signals’ correspondence to any class of activity largely depends on a particular expert, which can affect the study results. This issue is worth to highlight as application of ICA in EEG-studies become more and more popular. The ALICE toolbox is a special instrument to resolve these issues.

The developed web-application stores ICA data and makes it publicly available. This data includes annotations that are given by experts, which assign each component to the appropriate category. Moreover, the annotated dataset expands using the interface where each expert can make their own labeling. ALICE’s goal is to build a community where experts from the field of neuroscience, neurophysiology etc., share their ICA data and encourage each other to make the annotations. The low Kohen’s kappa coefficient and low inter-expert correlation in IC annotation obtained in our study points to high disagreement in components annotation evident even between two experts. Noteworthy, the only other crowdsourcing platform for IC classification (ICLabel, [12]) also report similar results: their mean inter-expert correlation was 0.50, ranging from 0.46-0.65, clearly pointing to different strategies of identification ICs. This finding emphasizes the need to study the reason for such low agreement between expert and to develop automatic IC classification toolbox that will work objectively.

The ALICE has a potential to unite the efforts of experts from different fields that are vital to develop an ML model that could be used in EEG studies for the objective assessment of various artifacts. Our baseline model is a clear evidence that the process of ICA artifacts selection can be easily automated using ML approaches. The novel aspects of the work include the algorithm for Mu and Alpha rhythm detection. The important point is that the model is publicly available and additionally can be used as a pre-trained model for posterior modifications for other tasks.

Subjective labeling and ML training was performed on a dataset of ICs obtained on EEG data recorded in pre-school and school age children, population with usually many artifacts and relatively unrepresented in the previous work on automatic IC extraction. The dataset consists of 630 ICA components acquired from 20 children, making up a unique publicly available dataset that can be used for various goals, e.g. for refitting new private models for ICA detection.

There are several points for future development of the project which are related to the annotation module and the ML module. The annotation module advances are related to the reorganization of available classes to mark into a hierarchical structure. People can first select the type of artefact and specify in more precisely, for example, Eyes-> Horizontal eye movements. Moreover, the first trial of expert annotations forces us to reestablish an expert policy and force them to choose no more than two types of IC classes to train our models using representative samples.

The ML module showed a high-performance rate for most classes. Although the Heart class was not detected, the reason for that is the lack of class representatives and a very low level of agreement between the annotators. Moreover, the results for Mu and Alpha rhythms and Eyes are also obtained with a low number of data samples. Nevertheless, the ALICE approach (including newly designed features for Mu and Alpha classes) showed good classification accuracy for ICs labeling even though the agreement between expert opinions was relatively poor. Still, for Heart and Channel Noise classes none of the trained models worked well. Probably new strong predictive features or more complex ML models (i.e. based on convolutional neural networks) should also be developed to address these types of artifacts. We compared our algorithms performance with results reported in other studies. In [12] authors report ROC curves with F1 scores. Eyes class F1 score is greater than 0.9, brain and muscles classes are in range between 0.8 and 0.9, which is higher than results obtained using our model; at the same time, the heart class, like in our case, is reported as uninformative. In [11] authors reported accuracy, sensitivity and false omission rates and provided full data for eye movements, eye blinks and muscle activity. The resulted F1 scores are greater than 0.9. In terms of our model the low agreement between experts that is came as an outcome of different labeling approaches might affect final score.

Current performance is based on two experts’ estimations, whereas a manifold of professional annotations produces more objective estimates for components labeling. In the future research, we aim to invite the wider expert community to label their datasets and expand current models’ abilities or future models to define functional nature of IC components.

To summarize, the main improvements implemented in ALICE as compared to previously developed toolboxes are the following:

- ALICE allows not only detection of noisy IC, but also automatic identifications of components related to the functional brain oscillations such as alpha and mu-rhythm
- the ALICE project accumulates different benchmarks based on crowdsourced visual labeling of ICs collected from publicly available and in-house EEG recordings, resulting into a constantly growing high quality IC dataset
- ALICE implements the new strategy for consentient labeling of ICs by several experts
- ALICE allows supervised ML model training and re-training on available data subsamples for better performance in specific tasks (i.e. movement artifact detection in healthy or autistic children)
- Finally, ALICE provides a platform for EEG artifact detection models comparison as well as a platform for neurophysiologists self-assessment based on established performance metrics.

Thus, a strength of the ALICE project implies the creation and constant updating of the IC database, which will be used for continuous improvement of ML algorithms for automatic labeling and extraction of non-brain signals from EEG. The developed toolbox will be available to the scientific community in the form of an online service and open-source codes.

